# Deformation-induced actuation of cells in asymmetric periodic flow fields

**DOI:** 10.1101/2021.09.30.462560

**Authors:** Sebastian W. Krauss, Pierre-Yves Gires, Matthias Weiss

## Abstract

Analyzing and sorting particles and/or biological cells in microfluidic devices is a topical problem in soft-matter and biomedical physics. An easy and rapid screening of the deformation of individual cells in constricted microfluidic channels allows, for example, the identification of sick or aberrant cells with altered mechanical properties, even in vast cell ensembles. The subsequently desired softness-specific segregation of cells is, however, still a major challenge. Moreover, aiming at an intrinsic and unsupervised approach raises a very general question: How can one achieve a softness-dependent net migration of particles in a microfluidic channel? Here we show that this is possible by exploiting a deformation-induced actuation of soft cells in asymmetric periodic flow fields in which rigid beads show a vanishing net drift.

---

The advent of microfluidic devices has fueled a multitude of experiments and applications, both in academia and industry [1–4]. In the biomedical context, high-throughput analyses of microfluidic droplets (‘microreactors’) or individual cells have gained particular interest [3, 5–7], not least as first steps towards future diagnostic usage. In this context, microfluidic devices for a rapid screening of individual cell stiffness [8, 9] are supposedly among the most prominent applications with a great potential for analyzing blood and biopsy samples: Exposing cells in narrow channel segments to strong Poiseuille flows, the resulting deformations can be evaluated by rapid imaging of the cells’ contour lines as they pass the constriction one by one within few milliseconds [8]. Using this approach, cells with aberrant mechanical properties, e.g. due to malignancy, sickle-cell disease, or malaria infections, can be detected even in complex cell mixtures like whole-blood samples [10].

Going beyond mere screening, key to further applications is a segregation of cells according to their mechanical properties. Using data from rapid imaging as an input for a downstream surface acoustic wave, cells have been shifted, for instance, into different arms of a y-shaped channel branching [11]. Yet, the need for high-speed (fluorescence) imaging with a sufficient quality, combined with an elaborate and sensitive feedback to address the acoustic wave, puts up the question whether a segregation and sorting of cells is also possible without any imaging by just exploiting the individual cell stiffness. This idea immediately raises a much more general and fundamental question: How can one achieve a softness-dependent net migration to differentiate particles with varying mechanical properties in a microfluidic channel?

Inspired by the principle of microscale ratchets that rely on spatially asymmetric periodic structures [12, 13] it has been hypothesized that exploiting the deformation of soft particles in temporally asymmetric periodic Poiseuille flows could induce a softness-dependent net migration [14]: Driving particles in a microfluidic channel at a forward flow rate *Q*_1_ for a time interval *τ*_1_, then reverting the flow and reducing the flow rate to *Q*_2_ for a time *τ*_2_ > *τ*_1_ leads to a restoration of the initial state of all fluid elements after a period *T* = *τ*_1_ + *τ*_2_ if *Q*_1_*τ*_1_ = *Q*_2_*τ*_2_. Since both, the flow rate and the slip velocity of a rigid sphere are proportional to the flow curvature, any residual drift due to Faxen’s law averages to zero over a period [15], i.e. non-deformable particles show no net drift after integer multiples of the period T. Soft particles, however, are deformed stronger during the first phase (0 < *t* < *τ*_1_) since *Q*_1_ > *Q*_2_, i.e. their effective extension perpendicular to the flow direction is reduced and hence they attain a higher effective drift velocity than rigid particles. In the restoration phase (*τ*_1_ < *t* < *T*) the deformation is less and the motion hence becomes more similar to rigid particles, eventually resulting in a net displacement Δ*x* per period *T*. Formulating this qualitative argument more thoroughly, the existence of such a softness-induced actuation was indeed confirmed via analytical calculations and computer simulations [14]. Yet, an experimental verification that soft cells can ratchet forward with each period, leaving stiff particles behind, as desired for a basic segregation mechanism, has been lacking so far.

Here we close this gap and confirm the prediction of a deformation-induced actuation of soft particles. In particular, we have quantified the migration of rigid plastic beads and soft red blood cells (RBCs) in a microfluidic channel design with an optimized hydraulic resistance to facilitate a rapid relaxation upon switching/inverting flow rates (see Materials and Methods in [16] for experimental details). As a result, we observed that beads did not show a significant net transport for integer multiples of the driving period *T* when applying symmetric or asymmetric periodic foward-backward flow profiles, whereas RBCs showed a marked actuation for an asymmetric driving. Our experiments therefore confirm the theoretically predicted possibility of a softness-induced segregation of particles, hence paving the way for a utilization of this effect in cell-sorting microfluidic devices.

As a first step of our experimental approach, we analyzed the shape of RBCs when subjecting them to different flow rates *Q* in a microfluidic channel (width: 10 *μ*m, height 100 *μ*m). Representative brightfield images of RBCs at flow rates *Q* = 800 *μ*l/h and *Q* = 200 *μ*l/h are shown in Fig. 1a,b. In line with previous observations [17, 18], RBCs assumed a parachute-like shape that was enhanced at higher flow rates. Extracting the contour lines of RBCs [16] allowed us to determine their projected area *A* and perimeter *ℓ*, with which we quantified the overall deformation via the deviation of the circularity from unity,

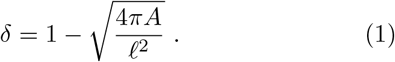

**FIG. 1:**
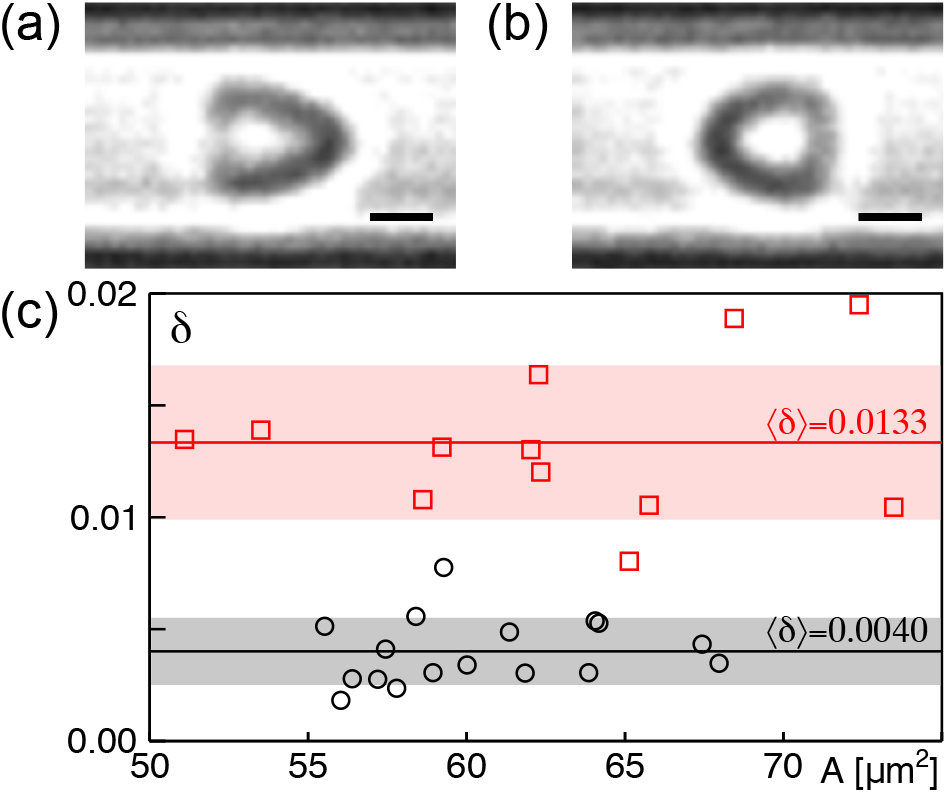
Representative brightfield images of RBCs in a microfluidic channel (width: 10 *μ*m, height 100 *μ*m) with flow rates (a) *Q*_1_ = 800 *μ*l/h and (b) *Q*_2_ = 200 *μ*l/h pointing to the right and left, respectively. Stronger deformations at higher flow rates are clearly visible; scale bar: 2 *μ*m. (c) Quantifying the cell deformation *δ* [Eq. (1)] confirms this visual impression: Deviations of individual RBCs from a circular shape (for which *δ* = 0) were significantly larger for *Q*_1_ (red squares) than for *Q*_2_ (black circles). Full lines and shaded regions indicate the respective mean and standard deviation.

This measure has also been used before for quantifying cell deformations in high-throughput experiments [8]. Confirming the visual impression, *δ* was significantly higher for *Q* = 800 *μ*l/h than for *Q* = 200 *μ*l/h (Fig. 2c). Hence, switching flow rates induces different shapes of RBCs, as required as an ingredient for the described ratcheting process.

**FIG. 2:**
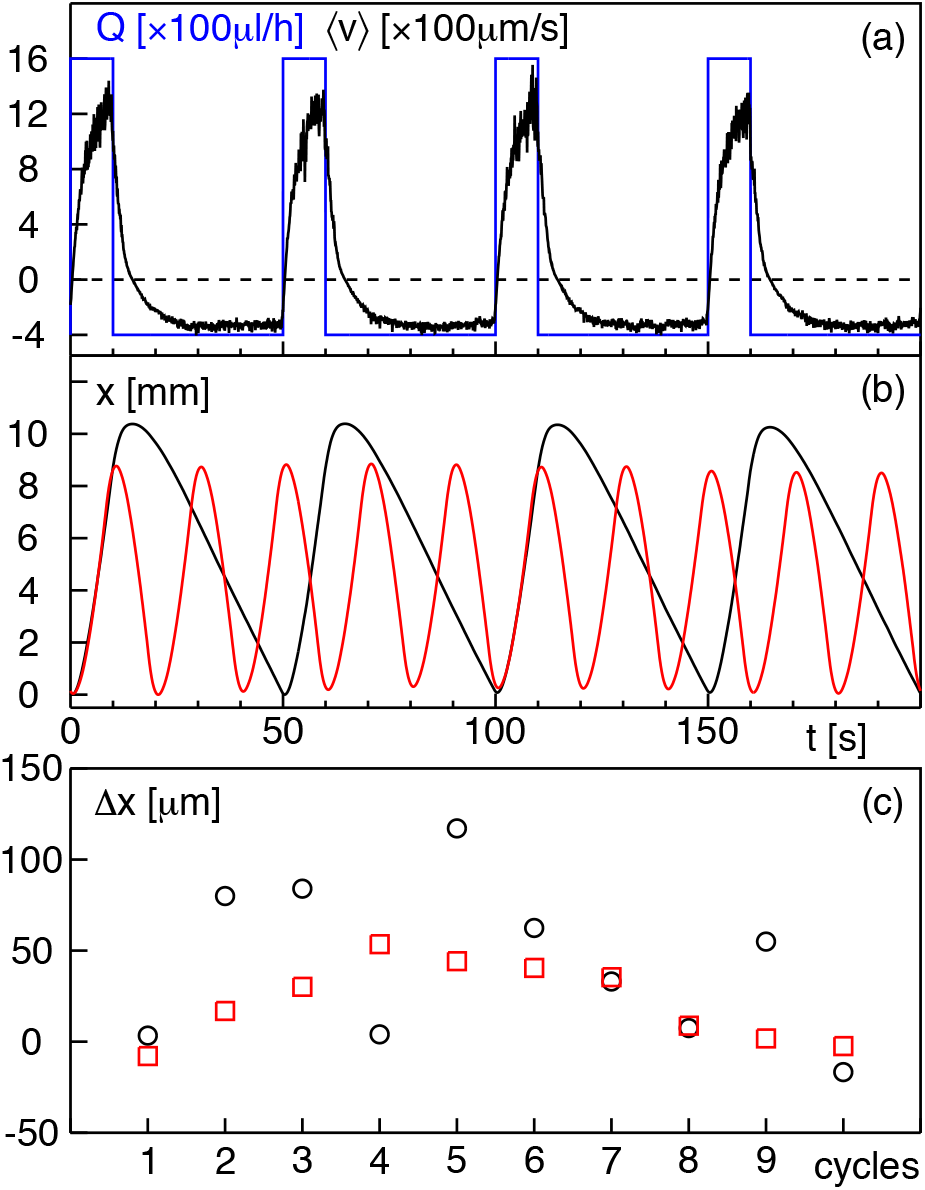
(a) Applying a periodic and asymmetrically switching dichotomous flow rate *Q*(*t*) in the microfluidic channel (blue line; foward: *Q*_1_ = 1600 *μ*l/h for *τ*_1_ = 10 s; backward: *Q*_2_ = 400 *μ*l/h for *τ*_2_ = 40 s) induces average maximum bead velocities that oscillate between plateau values 〈*v*〉 ≈ +1.4 mm/s and 〈*v*〉 ≈ −0.4 mm/s (black line). Please note that the periods *τ*_1_ and *τ*_2_ are sufficiently long to allow for a fair relaxation of the average velocity 〈*v*(*t*)〉 to the stationary value after reverting the flow. (b) Integrating 〈*v*(*t*)〉 over time yielded an average oscillatory excursion path *x*(*t*) for this asymmetric periodic flow rate (black line). For comparison, the path obtained for a symmetrically switching flow rate (*Q*_1_ = *Q*_2_ = 400 *μ*l/h and *τ*_1_ = *τ*_2_ = 10 s) is also shown (red line, multiplied by five for better visibility). In both cases, the minima in *x*(*t*) do not show signatures of a net drift. (c) In line with this observation, the deviation Δ*x* = *x*(*nT*) – *x*(0) from the initial position after *n* cycles is nonzero in both cases due to experimental imperfections, but no progressive increase (as expected for an actuation) is observed. This is in qualitative and quantitative contrast to soft RBCs (cf. Fig. 3c).

As a next step, we probed whether rigid beads show a net drift over several cycles when imposing an asymmetrically oscillating forward-backward flow field. Here, we concentrated on a forward flow of *Q*_1_ = 1600 *μ*l/h for *τ*_1_ = 10 s and a backward flow of *Q*_2_ = 400 *μ*l/h for *τ*_2_ = 40 s (see Fig. 2a), yielding a period of *T* = 50 s after which the initial state should be restored. The microscope’s field of view allowed for imaging 50-100 beads in every frame, situated at varying distances from the channel wall. Imaging and tracking these sets of beads at a frame rate of 140 Hz [16] allowed us to extract the beads’ average velocity 〈*v*(*t*)〉 (averaged over the parabolic flow profile sampled by the beads’ positions) as a function of time. Due to the finite number and length of tracks for each set of beads, the instantaneous average velocity showed some fluctuations, yet the temporal evolution of 〈*v*(*t*)〉 was clearly discernible (see Fig. 2a for a representative case). As expected for a channel with finite hydraulic resistance, every flow reversal required a finite relaxation time for approaching the asymptotic velocity associated with the new flow rate. Consequently, 〈*v*(*t*)〉 needed a few seconds after each switching of the flow before reaching a plateau value. Therefore, even when minimizing the hydraulic resistance, the possibility to reduce *τ*_1_ and *τ*_2_ to small values remains limited (see also discussion further below).

Integrating 〈*v*(*t*)〉 over time yielded the oscillating average position *x*(*t*) of the beads along the channel in response to the imposed flow field *Q(t)*. Results for an asymmetric and also a symmetric driving are shown in Fig. 2b. In both cases, the minima in *x*(*t*) did not show a progressive increase over several cycles, in contrast to what would be expected for an actuated motion. In accordance with this notion, quantifying the deviation from the inital position, after *n* periods, Δ*x* = *x*(*nT*) – *x*(0), did not show an increase for increasing number of cycles (Fig. 2c) albeit a nonzero value was observed due to experimental imperfections.

Performing the same tracking approach for beads and RBCs in the same system resulted in a marked difference between these two entities (Fig. 3): The integrated averaged position, *x*(*t*) showed a clear shift of the minima positions for soft RBCs whereas stiff beads returned to their original position (see magnification in Fig. 3a). Inspecting the associated profiles of the averaged velocity revealed that RBCs attained higher values for 〈*v*(*t*)〉 during the strong forward flow in the first phase (*nT* < *t* < *nT* + *τ*_1_). This discrepancy subsided for the softer backward flow during the second phase (*nT* + *τ*_1_ < *t* < (*n* + 1)*T*). The resulting net shift of an ensemble of RBCs over several periods of the oscillating flow, quantified via the minima positions *x*_min_ after integer multiples of T, was seen to grow linearly in time (Fig. 3c), in marked contrast to the restoring of bead positions (cf. Fig. 2c). Thus, soft RBCs showed a signficant net actuation in asymmetrically oscillating flows, as expected from theoretical considerations.

**FIG. 3:**
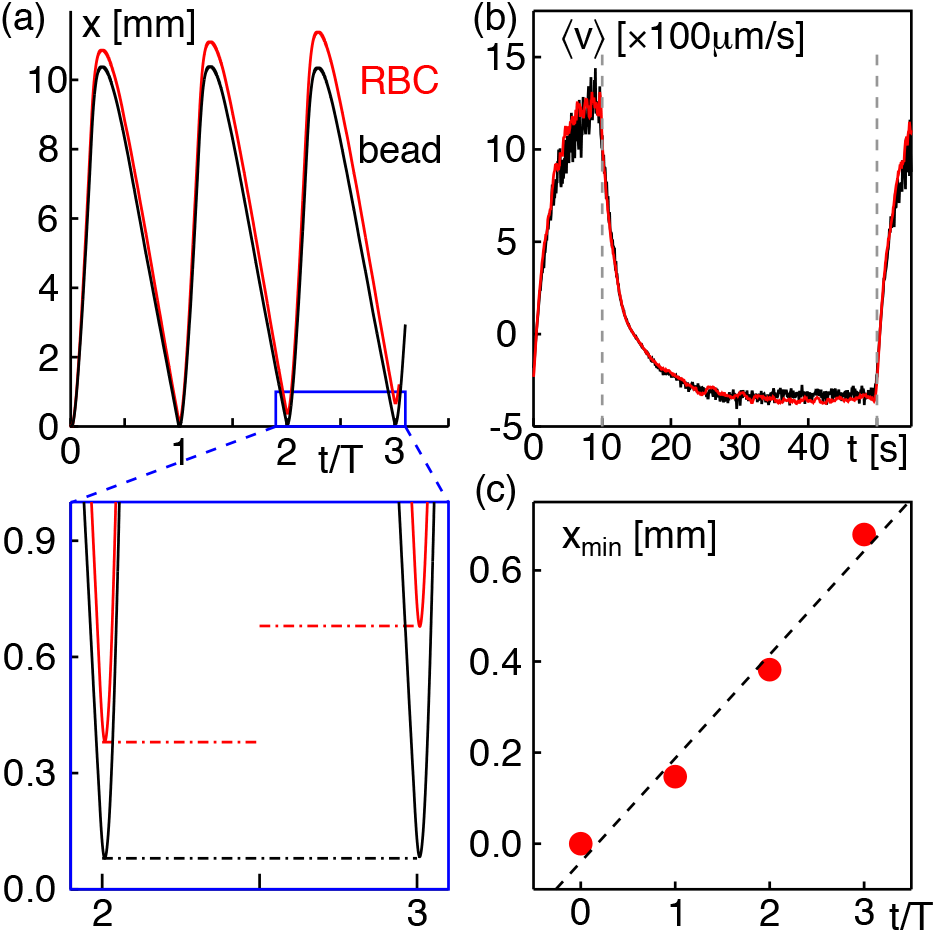
(a) Comparing the ensemble-averaged position *x*(*t*) of beads (black) and RBCs (red) in the same experiment reveals a marked progression of the minimum position for RBCs already after few periods *T* (highlighted by the magnification in the blue rectangular region; dash-dotted lines indicate the minima positions). (b) During the strong forward flow phase, RBCs attain a higher velocity than beads (red vs. black line), whereas the softer but longer backward flow during the second phase is very similar. Grey dashed lines indicate the times at which flow rates were reverted. (c) As a result, RBCs show a significant net actuation from the initial position for progressing periods, as quantified via the minima positions *x*_min_(*nT*) while beads remain basically at the inital position (cf. Fig. 2c). The dashed line is a linear guide to the eye.

So far, our evaluation approach relied on an ensemble average of many RBCs and beads. To complement these data, we also sought for a possibility to track individual RBCs. Since individual entities rapidly exit the microscope’s field of view due to the imposed flow rates, a direct long-term tracking was not possible. We therefore used an alternative approach: When analyzing the acquired image series, we measured the time shift Δ*T*(*n*) that was needed in addition to the integer number of periods nT to position the very same RBC into its original locus in the field of view. Using the average velocity 〈*v*(*t*)〉 determined from the collection of beads (cf. Fig. 2a), this time shift was then translated into a traveled distance *ξ* that represents the net shift of this particular RBC along the channel. Due to the actuation, a relation *ξ* ~ Δ*T*(*n*) ~ *nT* is expected and hence a linear increase of *ξ* with time is anticipated. Inspecting the travel distance for individual RBCs (see Fig. 4 for representative examples) confirmed this very expectation, in favorable agreement with the aforementioned finding of an average net actuation that grows linearly with the number of cycles (cf. Fig. 3c).

**FIG. 4:**
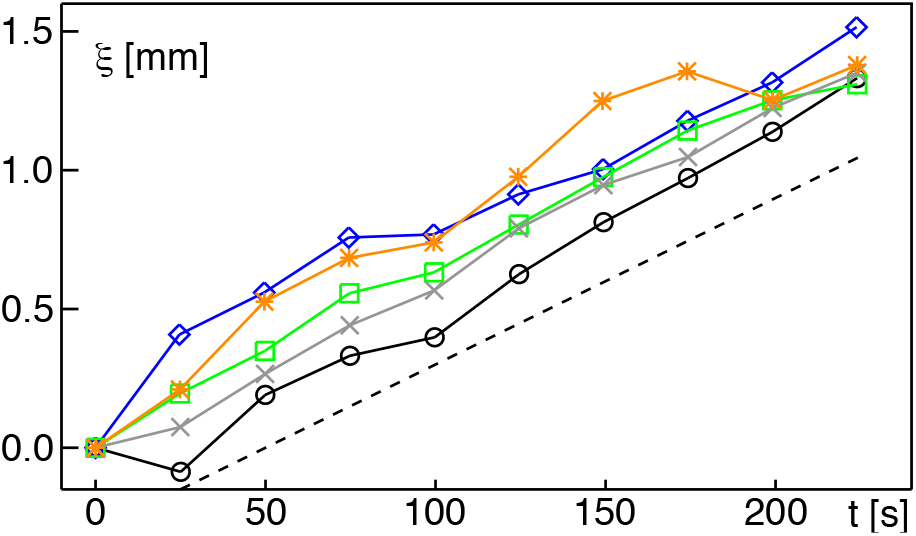
Tracking individual RBCs (see main text and [16] for details) reveals a net actuation *ξ* that grows linearly with the number of periods, in agreement with the average drift of an ensemble of RBCs displayed in Fig. 3c.

In summary, we have demonstrated here that subjecting soft RBCs to asymmetric oscillatory flow fields in a microfluidic channel can induce a net actuation that grows linearly with the number of driving periods, whereas stiff beads essentially show no net transport. This finding is in agreement with theoretical considerations [14] and provides a basic unsupervised mechanism to segregate particles according to their softness. Starting from our proof-of-principle data, one may envisage a usage of the effect, e.g. in biomedical applications that require cell sorting according to stiffness. While the finite relaxation times due to the hydraulic resistance certainly will hamper a high-throughput scheme for individual cells, the approach shown here is nicely suited for performing the segregation of large cell ensembles within a reasonable time. It may therefore be used for a sorting of complex mixtures after (alternatively: prior to) screening individual cells in high-throughput devices described earlier [8, 9]. Since the scheme that we have presented here only relies on the flow control device (e.g. syringe pumps), which is typically already part of an existing microfluidic platform, no additional equipment (such as a high speed camera and a surface acoustic field emitter) is required. It therefore extends the toolbox of microfluidics-based techniques with an efficient and inexpensive approach.

## Acknowledgments

We thank W. Zimmermann and W. Schmidt for suggesting the experiments. Financial support by the VolkswagenStiftung (Az. 92738) and by the EliteNetwork of Bavaria (Study Program Biological Physics) is gratefully acknowledged.

## I. MATERIALS AND METHODS

### A. Design and production of microfluidic channels

To probe a net actuation of soft particles in an (asymmetrically) oscillating Poiseuille flow, we designed a microfluidic channel with a height of 100 *μ*m, a width of 10 *μ*m, and a total length of 45 mm (see Fig. S1). To reduce the relaxation times after switching the flow rate, a bypass channel with a width of 500 *μ*m was added in parallel to the probe channel. Due to the similar hydraulic compliance of the bypass channel and the inlet, an optimal reduction of the relaxation times was achieved. RBCs and beads were fed into the device via a third inlet close and perpendicular to the probe channel (see Fig. S1).

**FIG. S1.**
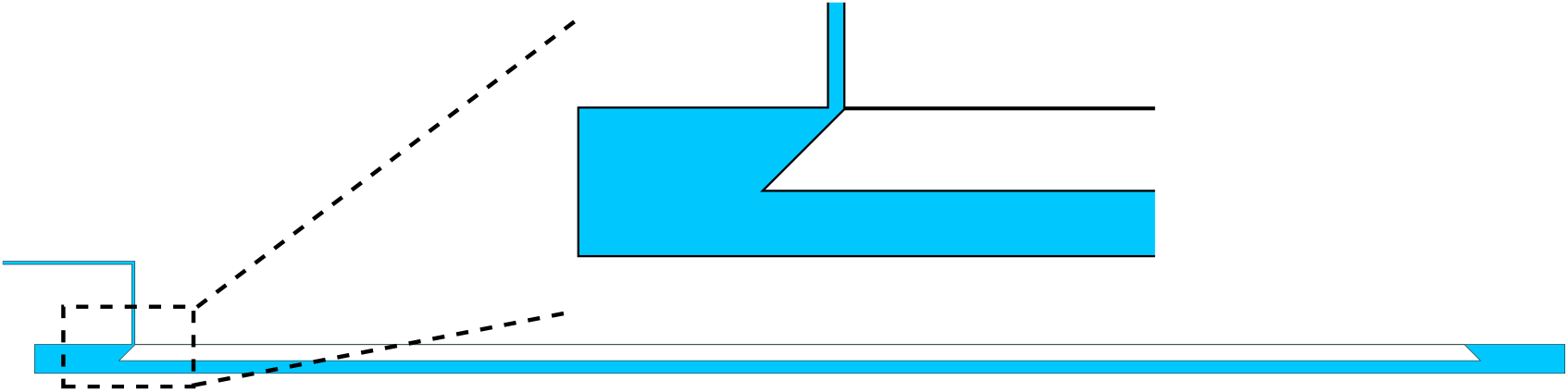
Scheme of the microfluidic device with the magnification highlighting the two parallel channels where the wider one was introduced to reduce relaxation times upon switching flow rates. A third, perpendicular inlet was used to introduce beads and cells.

Microchannels were produced by standard soft lithography. The master mold was manufactured using a negative photoresist (SU-8 2050, microchem) spread to a height of 100 *μ*m by spin coating on a silicon waver. After soft-baking on two hotplates (65°C for 5 min and 95°C for 20 min) the channel design was directly written with a microwriter (MicroWriter ML3 Baby+, Durham Magneto Optics). After the post exposure bake (65°C for 5 min and 95°C for 10 min), unexposed photoresist was removed by successively washing the wafer with developer (mr-Dev 600) and isopropanol, followed by drying with compressed air. Polydimethylsiloxan (PDMS) based microfluidic chips were then drawn from the master. To this end, PDMS (Sylgard™ elastomer 184, DOWSIL) was mixed with curing agent in a 10:1 ratio and poured onto the wafer after 2 min. Degasing was performed two times for 2 min in a Duran desiccator (Schott, Germany). After baking in the oven at 75°C for 3.5 h, the cured elastomer was removed from the wafer with a scalpel. Inlets and outlets were punched into the PDMS with a stainless steel puncher (Harris Uni-Core, 1.2 mm). The device was bonded to a microscope glass slide (cleaned by sonication in isopropanol for 10 min and dried in nitrogen flow) following a plasma exposure for 20 s at 122 Pa. Bonding was strenghtened by storage at 75°C in the oven for 45 min. The chip was soaked in Milli-Q water overnight prior to the experiment for equilibration.

Flows in these devices were driven by syringe pumps (CETONI neMESYS low pressure module V2, controlled by base module BASE 120). Connections between syringes and chip were done via PTFE teflon tubings with an inner 2 diameter of 0.5 mm and an outer diameter of 1.58 mm. To eliminate flows introduced by a loose connection between chip and tube, tubings were glued to the inlets by a two component epoxy adhesive. Opposing flows have been applied to the inlet and the outlet.

### B. Sample preparation

Venous blood was donated by the first author in accordance with all health and safety regulations. Fresh blood was contained in EDTA tubes (S-Monovette) and the subsequent isolation of red blood cells (RBCs) was performed as described before (Flormann *et al*., 2017). In brief, RBCs were pelletized by centrifugation (700g for 3 min) using a Heraeus Fresco 17 centrifuge. The supernatant was removed and the cell pellet was washed three times with PBS. RBCs were resuspended by gentle repetitive pipetting of the pellet. To minimize osmotic pressure on the cells, the RBC medium was tuned to an isotonic condition via phosphate-buffered saline (PBS) (Fischer, 2010). For convenience, the viscosity of the medium was adjusted to the cellular viscosity via 12.5 w% dextran (molecular weight 70 kDa) (Carrasco *et al*., 1989). All experiments were carried out within four hours after blood donation. As tracer beads, 2 *μ*m polystyrene-particles (micromer, 01-02-203) with a carboxylic acid functionalized surface were used at a concentration of 1 mg/ml. RBCs in the final solution have been diluted 200× compared to the pellet concentration.

### C. Imaging and tracking

Brightfield imaging of the sample channel was performed with an inverted microscope (ZEISS Axiovert 25) with a 20x objective (Zeiss Achroplan 20x/0.45 Ph2 440141) and a 6 W halogen lamp for illumination. The field of view was 490 *μ*m×40*μ*m; size calibration was performed with a microscopy scale bar (0 – 100 *μ*m). Images were acquired at a speed of 140 frames per second via a sCMOS camera (Manta G-201B, ASG Allied Vision GmbH).

To analyze their displacement, beads were automatically tracked using a linear motion LAP tracker within FIJI/Trackmate (Tinevez *et al*., 2017). Time and position of each bead were subsequently extracted by a custom-made Matlab code. RBCs have been tracked manually to prevent ambiguities in particle identification due to a high bead concentration. Based on the positions between successive frames, the average velocities of beads and RBCs was calculated for each frame. Complementary to the average ensemble dynamics, the shift of individual cells over multiple periods was analyzed by extending the reverted flow, *Q*_2_, for an additional time ΔT that was required to restore the RBCs initial position. Using this approach over many cycles, the total shift of individual RBCs over time became accessible (see Fig. 4 of the main text for examples).

### D. Contour extraction of cells

A set of representative RBCs was first manually selected for further quantification. The subsequent workflow for contour detection of these RBCs, using Matlab, was as follows:

1. Manual cropping of a rectangular region of interest around a RBC, yielding the initial image *I*_ini_, see Fig. S2a.
2. Binarization of *I*_ini_, yielding *I*_th_, via an adaptative thresholding (Matlab function *adaptthresh*), followed by a hole-filling step (Matlab function *imfill*) to yield *I*_th2_, see Fig. S2b.
3. Distance transform of *I*_th2_ (Matlab function *bwdist*) yielding an image *D*, see Fig. S2c with the greyscale bar indicating the pixel distance to the closest dark pixel in *I*_th2_.
4. Watershed transform of –*D* (Matlab function *watershed*) yielding an image *W*, see Fig. S2d.
5. Masking of W by *I*_th2_, yielding an image *W_m_*, see Fig. S2e.
6. Extraction of the connected component with centroid closest to the image center. The resulting contour line *C* is superimposed to *I*_ini_; in Fig. S2f.
7. Active contour evolution of C based on the gray scale values of *I*_ini_; as described in (Chan and Vese, 2001) (Matlab implementation in function *activecontour*, Chan-Vese method, 400 iterations, ‘SmoothFactor’ set to four and no contraction bias). The resulting contour, *C*_ac_, is shown in Fig. S2g.
8. Dilation of *C*_ac_ with a disk as structural element. The best fit with a visual estimate of the RBC contour was found with the largest disk radius so that 〈*I*_corona_〉/(〈*I*_mask_〉 > 0.1. Here, the index ‘mask’ refers to pixels within the initial contour, the index ‘corona’ denotes pixels in the surrounding after contour dilation, see Fig. S2h. The resulting contour, *C_d_* is shown in Fig. S2i.
9. Shape filtering. The filtered contour consists of the Fourier components of *C_d_* including terms up to third order, see Fig. S2j.

**FIG. S2.**
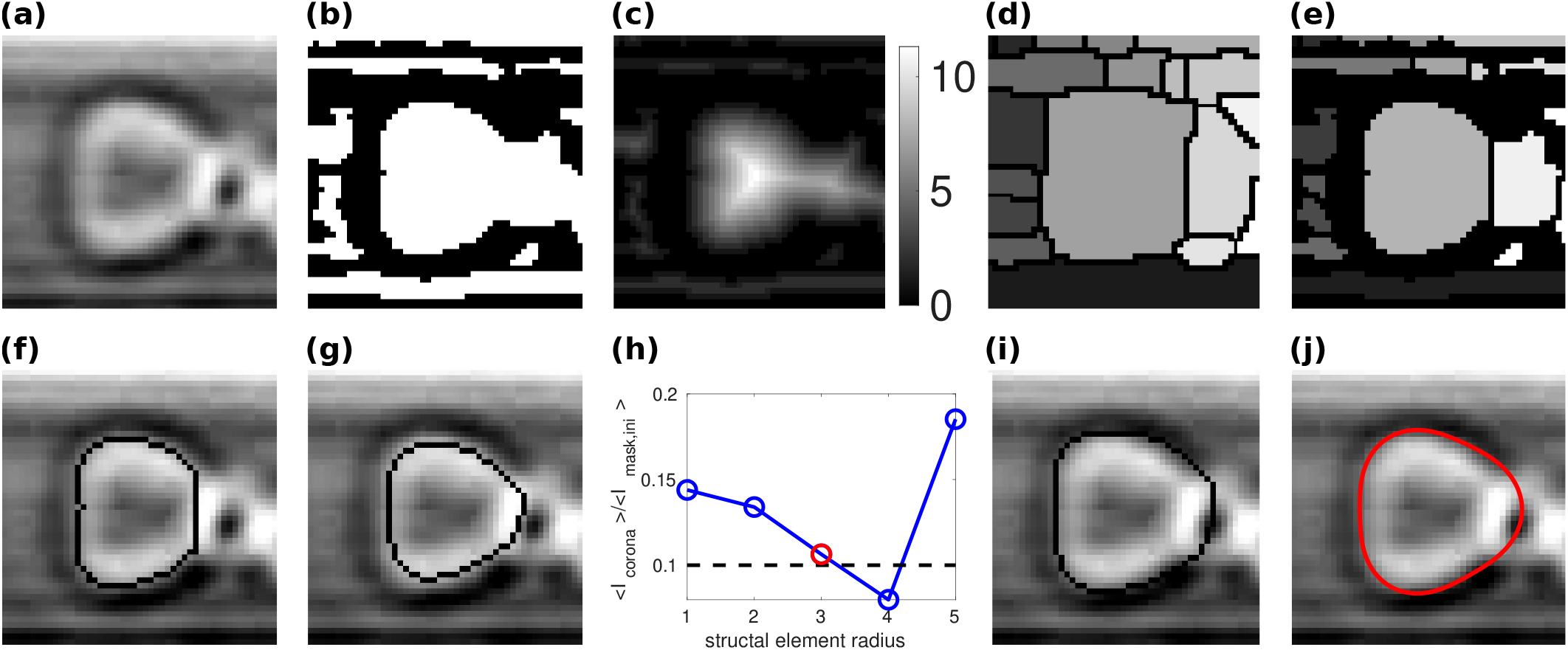
Automatic contour detection of a RBC as described in the list of actions. (a) Initial image. (b) Automated thresholding. (c) Distance transform. (d) Watershed transform. (e) Masked watershed transforn by (b). (f) Resulting contour. (h) Optimal dilation radius search: optimum found here for a structural element being a disk of radius 3 px. (i) Contour after dilation. (j) Contour after Fourier smoothing up to the third harmonic included.

Representative contour lines for RBCs subjected to different flow rates, as obtained via the outlined approach, are shown in Fig. S3.

**FIG. S3.**
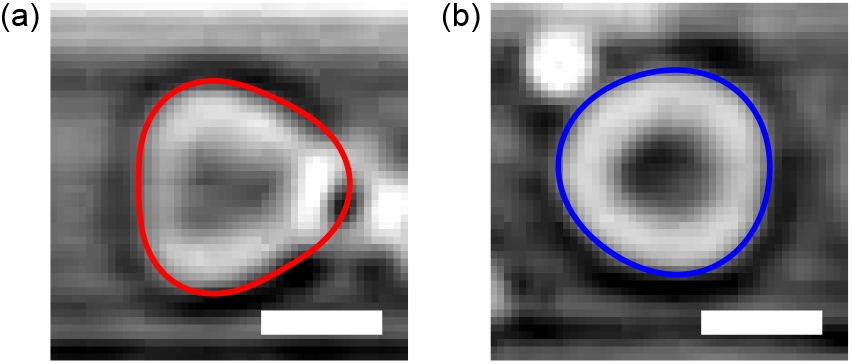
Examples of RBCs and their extracted contour lines for (a) fast and (b) slow flow rates; scale bar: 5 *μ*m.

